# Establishment and Evaluation of Rat Models of Cervical Spondylopathy with Yin Deficiency Syndrome

**DOI:** 10.1101/2023.04.06.535905

**Authors:** Wenlong Yang, Muqing Liu, Qinran Sun, Lei Liu, Wenqing Wu, Peng Gao, Fangming Liu, Zhizhen Liu

**Affiliations:** Department of Rehabilitation Medicine, The First Affiliated Hospital of Shandong First Medical University & Shandong Provincial Qianfoshan Hospital, Shandong Institute of Anesthesia and Respiratory Critical Medicine; School of Acupuncture-Tuina, Shandong University of Traditional Chinese Medicine, Shandong province, China

## Abstract

**Objective:** Cervical spondylosis (CS) with Yin deficiency syndrome is an important classification of traditional Chinese medicine (TCM) symptoms of CS. However, there is no animal model for studying this TCM syndrome. This study aimes to establish and evaluate rat models of cervical spondylopathy with Yin deficiency syndrome.

**Method:** Thirty-six Sprague–Dawley male rats were randomly divided into the blank control (control), CS and CS with Yin deficiency syndrome (YCS) groups (n = 12 rats/group). CS was induced using cervical static–dynamic imbalance to mimic disk degeneration (except in the control group). After 30 days of the CS rat model, rats in the YCS group were subjected to sustained sleep deprivation for 168h. After different induction times, the rats in each group were observed for behavior and weight. Pain behavior was assessed by a withdrawal response to von Frey filament application, and heart rate and blood pressure were measured using a rat noninvasive sphygmomanometer. Intervertebral disc pathology sections were observed using hematoxylin and eosin staining and an electron microscope. Western blotting was used to evaluate the protein level of collagen-II, Bcl-2, Bax and Bcl-2/Bax expression in the cervical intervertebral disc. Determination of related laboratory serum indexes, cyclic adenosine monophosphate, and cyclic guanosine phosphate were conducted using enzyme-linked immunosorbent assay.

**Results:** The laboratory indexes in the YCS group were significantly different from those in the control and CS groups (P < 0.05), and indicators of 72-, 120-, and 178-h sleep deprivation showed varying degrees of difference from those of the CS group.

**Conclusion:** After establishing a model of CS, continuous sleep deprivation for 72 h was used to create a rat model of CS with Yin deficiency syndrome. The established rat models of CS with Yin deficiency syndrome met the clinical and Chinese medicine characteristics, and thus, they can be expected to become an ideal model for studying CS with Yin deficiency syndrome in the future.

## Introduction

Cervical spondylosis (CS) is a cervical intervertebral disc degenerative change, and its adjacent structure of secondary pathological changes involves the organizational structure [1]. It is a common public health concern, accounting for a large proportion of frequently occurring occupational and disabling factors [2]. Traditional Chinese medicine (TCM) has a definite curative effect and unique advantages in the treatment of CS [3], but there are few studies on the TCM syndrome model of CS. Therefore, establishing an animal model with the characteristics of the TCM syndrome is an important task for the adaption to the development of medicine.

A previous study has shown that some patients with CS have increased pain in the neck, shoulder, arm, and other parts at night that makes them unable to sleep, and these are accompanied by palpitation, irritability, and slight fever in the afternoon, night sweats, insomnia, and dreaminess [4]. TCM believes that long-term fatigue injury of the cervical spine consumes Yin in the body. When a patient is in a state of Yin deficiency for an extended period, the symptoms of Yin–Yang metabolism disorder appear in the meridians, viscera, and the whole body [5], causing CS and the above symptoms [6]. At present, there are neither reports, nor studies on animal models of CS with Yin deficiency syndrome, thereby hindering the development of experimental research on this disease. Thus, in this study, we aimed to establish rat models presenting the characteristics of CS with Yin deficiency syndrome. Cervical static–dynamic imbalance and sustained sleep deprivation were used to induce CS with Yin deficiency syndrome in rats.

## Materials and Methods

### Animals and models

#### 1. Animals

Thirty-six male Sprague–Dawley rats weighing 200 ± 20 g were provided by Jinan Pengyue Experimental Animal Breeding Co., Ltdr (SCXK Lu 2019–0003, Shandong, China). The animals were maintained under a standard temperature of 23 °C ± 2 °C and a 12-h light–dark cycle. The rats were grouped into three rats per cage with free access to standard diet and water. All procedures were reviewed and approved by the Ethics Committee of the First Affiliated Hospital of Shandong First Medical University & Shandong Provincial Qianfoshan Hospital ([2019]no. S177).

#### 2. Groups

All rats were randomly divided into three groups with 12 rats in each: blank control group (control group), CS group (CS group), and CS with Yin deficiency syndrome group (YCS group).

#### 3. CS models

Twenty-four rats were randomly selected to construct models of CS according to the reference and preliminary study [7]. After 1 week of adaptive feeding, 24 rats were randomly selected to be anesthetized by intraperitoneal injection of 3% pentobarbitone sodium at a dosage of 30 mg/kg. The anesthetized rats were put on a fixed plate in a prone position. After sterilizing the skin with anerdian, a posterior cervical median incision was made to expose the tissue from the atlanto-occipital joint to the second thoracic vertebrae. The fascia, platysma muscle, cervical trapezius, and rhomboideus were then separated, and the bilateral muscular area was blunt stripped until the deep layer of the muscle was exposed. A cut was made across the splenius cervicis, head, cervical, musculus longissimus atlantis, costo cervicalis, and musculus semispinalis capitis to fully expose the cervical vertebrae, and the C2–C7 supraspinous and interspinous ligaments were snipped. After the bleeding stopped, the bilateral sacrospinal muscles and skin were sutured. All rats were injected with penicillin for 3 days after the operation to avoid infection. The stitches were not removed. Two rats were selected and sacrificed from the model group 30 days after modeling. The C4/5 and C5/6 intertebral disks were taken and fixed.

Hematoxylin and eosin (H&E)-stained sections were used to observe the histomorphology of the cervical intervertebral disc under a light microscope. We observed that the morphology and structure of the intervertebral disc in the control group were normal, but the model group exhibited morphological changes, disorder of annulus fibrosus, and shrinkage of the nucleus pulposus (Figure 1). Meanwhile, After staining with uranyl acetate and lead citrate, the ultrastructure of the nucleus pulposus was observed under an electron microscope. The cells in the control group were round and oval. Th cytomembrane was visualized, and the nucleus was in the center with a smooth and integral nuclear membrane. The cells in the CS group were irregular in shape, and the nuclear membrane structure was uneven (Figure 2). Finally, in the model group, the neck-skin response thresholds to the mechanical stimuli applied by von Frey filaments were significantly decreased 30 days after surgery (Figure 3a). The CS model was successfully established.

**Figure 1.**
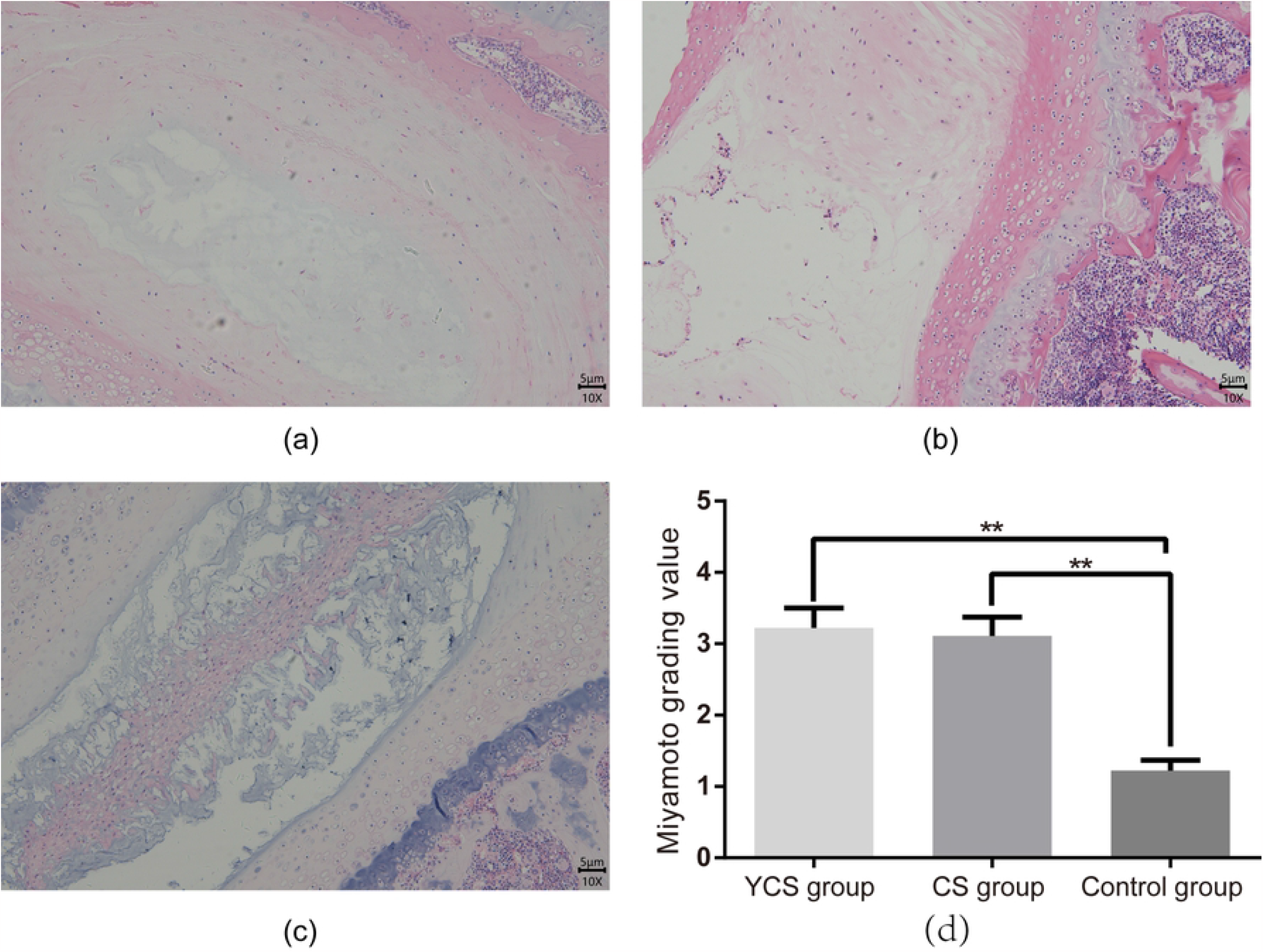
H&E staining and scoring for cervical intervertebral disc. The representative H&E staining images of cervical intervertebral disc of rats from. (a) YCS group (after 168 h of sleep deprivation), (b) CS group, (c) Control group, (d)The scoring analysis for cervical intervertebral disc based on H&E staining. Data are represented as mean ± SEM (n = 9), scale bar = 5 μm, \*\**P* < 0.01.

**Figure 2.**
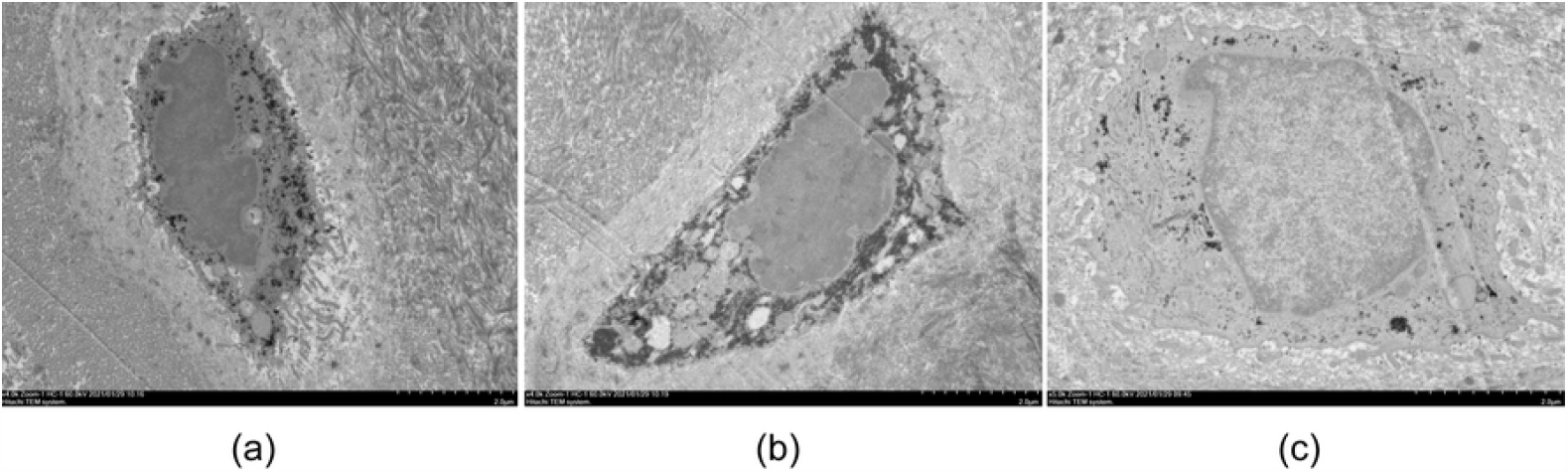
The ultra-structure of the nucleus pulposus in different groups under transmission electron microscope. Representative images for ultra-structure of the nucleus pulposus in different groups under transmission electron microscope. (a) YCS group (after 168 h of sleep deprivation), (b) CS group, (c) Control group. Note: scale bar = 2 μm.

**Figure 3.**
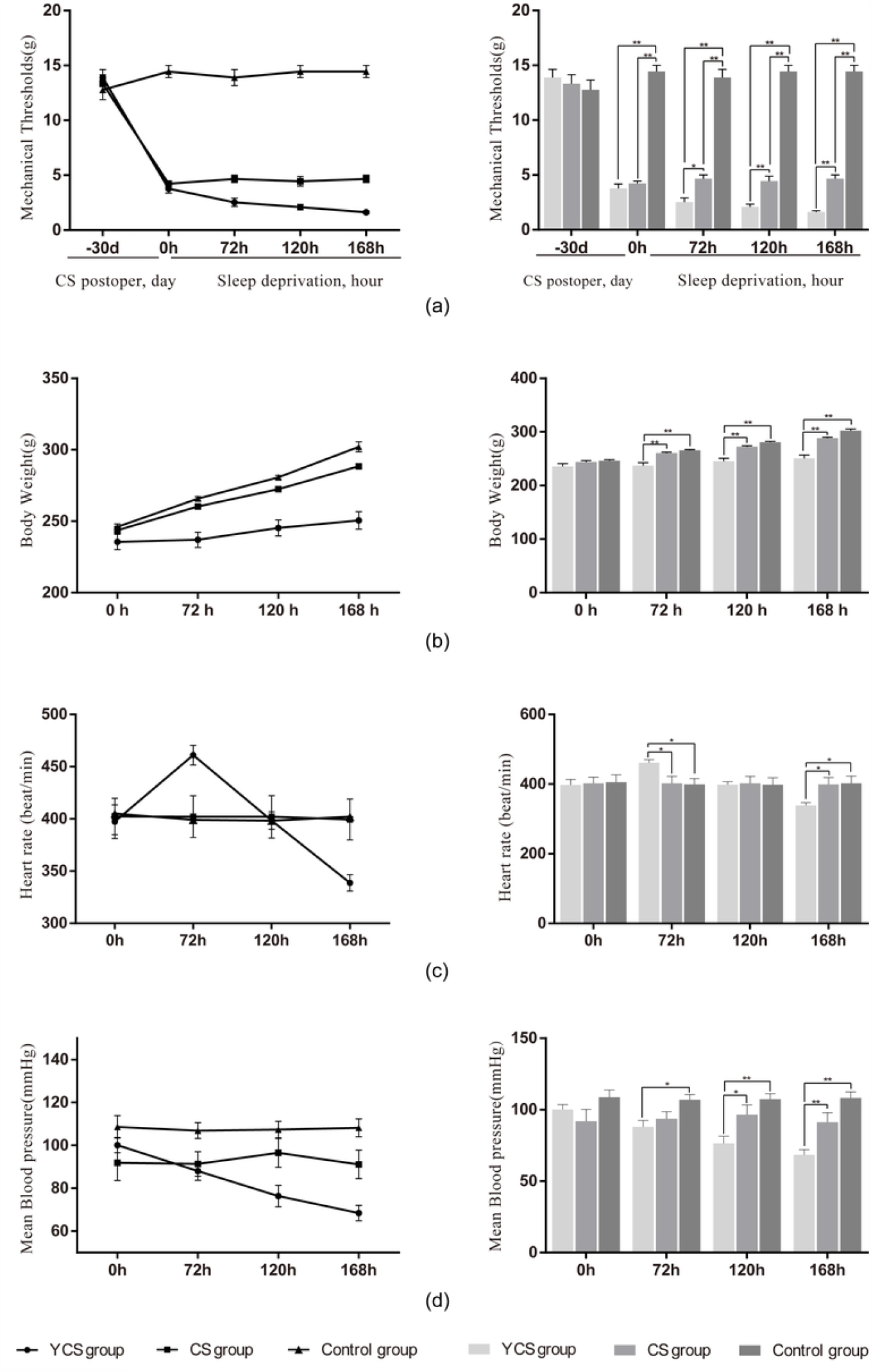
Indexes of Yin deficiency Syndrome in Animal Models. (a) Mechanical sensitivity of each group measured for 30th day after surgery and 168 hours of sleep deprivation. (b) Body weights of each group measured for 168 hours of sleep deprivation. (c) Heart rate of each group measured for 168 hours of sleep deprivation. (d) Mean blood pressure of each group measured for 168 hours of sleep deprivation. Data are represented as mean ± SEM (n = 9), \**P* < 0.05, \*\**P* < 0.01.

#### 4. YCS models

After the successful manufacture of the CS model, the rats were continuously deprived of sleep for 168 consecutive hours. The behavioral changes, weight changes, pain threshold changes, heart rate and blood pressure changes of the three groups were observed and recorded.

#### 5. Sleep deprivation

To induce total sleep deprivation (TSD), we used an automatic TSD apparatus (XR-XS107, Xinruan Co.Shanghai, China). The apparatus was made of clear Plexiglas (464 × 300 × 180 mm). A linearly cycling agitator was placed at the bottom of the apparatus, which physically disturbed the rats, thus preventing sleep. Through magnetism, the platform pulls the agitator along the cage floor. The agitator moves in alternating directions in a linear path along the length of the apparatus. In the current experiments, the speed was set at approximately 25.4 mm/s to provide food and water to the rats. The other two groups were placed in a narrow space similar to the rats in the sleep deprivation instrument, but they were not deprived of sleep.

#### 6. Behavioral observation

Experimenters blinded to the group allocation conducted the behavioral observation. Behavior related to the Yin deficiency syndrome in the rats was observed and recorded before sleep deprivation (hour 0) and at 72, 120, and 168 h after sleep deprivation. The observation contents included weight, mental state, fur color, caught resisting, stool, and urination [8].

#### 7. Von Frey test for mechanical allodynia

Pain-related behavior in rats was tested before surgery (day 0) and 30 days after surgery (before sleep deprivation) and at 72, 120, and 168 h after sleep deprivation. All experiments were carried out in a quiet room under soft light between 9 a.m. and 6 p.m., and the room temperature was maintained at 25 °C throughout behavioral testing. The hairs in the neck and back measured regions of the mice were removed using a hair clipper (HC1066, Philips, Netherlands). The rats were then habituated to a box made of black wire mesh for 30 min or until cage exploration and major grooming activities ceased. Withdrawal responses to mechanical stimulation were determined using calibrated von Frey filaments applied from above the cage to the skin 2 mm adjacent to the wound and the same area on the noninjured neck skin of the control rats. We set the cutoff value at 15 g, and the first filament in the series that evoked at least three responses (from the five applications) was regarded as the threshold. If there was no response to the cutoff force, 15 g was recorded. A quick withdrawal of the neck after the filament became bent was defined as a response [9].

#### 8. Heart rate and blood pressure measurements

Heart rate and blood pressure in rats were tested before surgery (day 0) and 30 days after surgery (before sleep deprivation) and then at 72, 120, and 168 h after sleep deprivation. All experiments were carried out in a quiet room under soft light between 9 a.m. and 6 p.m., and the room temperature was maintained at 25 °C throughout behavioral testing. After rats were calm, heart rate and blood pressure were evaluated using the tail-cuff method with a noninvasive automatic blood pressure recorder (BP-2010A, Ruanlong Co., Beijing, China). Each value was the average of at least three consecutive measurements [10].

#### 9. Specimen collection

The experimental period of sleep deprivation was 168 h. During the experiment, 2 mL of blood was extracted from the femoral vein and injected into a coagulant test tube before sleep deprivation and at 72, 120, and 168 h. The blood sample was allowed to stand at 4 °C for 2 h and then centrifuged in a cryocentrifuge at 4 °C and 5000 rpm/min for 25 min. The supernatant was then collected and stored at a temperature of -80 °C until enzyme-linked immunosorbent assay (ELISA). After the rats were killed, the specimens of rat C5/6 cervical intervertebral discs were collected and fixed in 4% paraformaldehyde solution and 2.5% glutaraldehyde for electron microscope and histology analysis. The disk tissues of C4/5 were collected and stored at -80 °C for western blot test.

#### 10. ELISA

We used ELISA to detect the concentration of cyclic adenosine monophosphate (cAMP) and cyclic guanosine monophosphate (cGMP) in serum. The specific operations were carried out according to the instructions of each kit, taking cAMP as an example. The serum samples were melted at room temperature, and the supernatant and cAMP standard were dispensed into the wells of an ELISA plate. We dispensed 50 μL of a biotinylated detection antibody diluent, sealed the wells, and incubated the plate at 37 °C for 45 min. Next, we discarded the solution in the wells and washed the wells with wash solution three times. We then dispensed 100 μL of concentrated horseradish peroxidase (HRP) -conjugated antibody solution into the wells, sealed the wells, and incubated the plate at 37 °C for 30 min. Next, we discarded the solution in the wells and washed the wells five times with wash solution. Finally, we added 90 μL of substrate reagent to each well, gently mixed and incubated for 15 min at 37 °C before adding 50 μL of stop solution to each well. We then read the optical density of each well at 450 nm (E-EL-0056c, Elabscience Biotechnology Co., Wuhan, China).

#### 11. H&E staining

The intertebral disks of the rats were fixed with 10% formalin and decalcified for 2 weeks by EDTA. Paraffin sections were deparaffinized, rehydrated, and then counterstained with hematoxylin. Finally, morphological changes were observed under a microscope. An intervertebral disc was evaluated using the grading standard as previously reported [7]. Morphological changes were observed under a microscope.

#### 12. Ultrastructure examination under an electron microscope

The intertebral disk without vertebral body was taken and subjected to the following method of sample preparation: rinsing, fixation, embedding, penetration, and sectioning. After staining with uranyl acetate and lead citrate, the ultrastructure of the nucleus pulposus was observed under an electron microscope.

#### 13. Western blot

We extracted the total proteins of the intervertebral disc tissues with ice-cold RIPA buffer containing phenylmethanesulfonyl fluoride (1 mM). The samples were then centrifuged at 12,000 rpm for 20 min. We used the BCA Protein Assay Kit (P0012, Beyotime Biotechnology, Co., Shanghai, China) to measure the protein concentration in the supernatant. The samples were boiled with protein loading buffer, and 30 μg of total protein was separated by sodium dodecyl sulfate-polyacrylamide gel electrophoresis and then transferred onto polyvinylidene fluoride (PVDF) membranes. The PVDF membranes were blocked with 5% skim milk in TBST (20 mM Tris-HCl, pH 7.5, 150 mM NaCl, and 0.05% Tween-20) for 2 h at room temperature. The membranes were incubated with primary antibodies against anti-collagen-II (1:1000 dilution, ab188570, abcam Co., Cambridge, UK), anti-Bcl-2 antibody (1:500 dilution, ab196495, abcam Co., Cambridge, UK), anti-Bax antibody (1:1000 dilution, ab32503, abcam Co., Cambridge, UK), and GAPDH (1:2,000 dilution, E-AB-20059, Elabscience, Wuhan, China) at 4 °C overnight. The PVDF membranes were washed with TBST and incubated with HRP-conjugated goat antirabbit immunoglobulin G (H + L) secondary antibody (1:5000 dilution, EF0002, sparkjade Co., Shandong, China) for 2 h at room temperature. Protein bands were detected using the enhanced chemiluminescence method. The band intensity was quantified using Image J software (public domain), and the mean gray value of each band was used for further statistical analysis.

#### 14. Statistical method

We analyzed the data using SPSS 23.0 software (IBM Corporation, Armonk, NY) for statistical analysis and GraphPad Prism 8.0.1 (San Diego, CA, USA) to prepare the graphs. All data are expressed as mean ± standard error of the mean(SEM). Unpaired Student t test was used to compare two groups. One-way analysis of variance (ANOVA) or two-way repeated-measures ANOVA, followed by post hoc Tukey test, was used for multiple comparisons of normally distributed data. Nonnormally distributed data were analyzed using Kruskal–Wallis nonparametric analysis. A P value < 0.05 was considered statistically significant.

## Results

### Indexes of Yin deficiency syndrome in animal models

#### 1. Behavioral characteristics of the animal models

We observed the behavioral characteristics of rats subjected to 0 to 168 h of sleep deprivation. The YCS group before the sleep deprivation intervention was mentally active; their fur was moist and white; their lips, nose, and tail were ruddy; caught resisting was moderate; urination was normal; and stools were dry and formed. After 72 h of sleep deprivation, the YCS group showed certain excitability; increased exploration behavior; sensitivity to external sound, light, and other stimuli; slightly dull fur; red and glossy lips, nose, claws, and tail; intense caught resisting; biting; yellow urine with a peculiar smell; and dry stool. After 120 h of sleep deprivation, the rats were mentally tired; their activity decreased; their fur was slightly haggard and fluffy, white, and slightly yellowish; the color of the lips, nose, and tail was dim; caught resisting was light; they had oliguria; and their stool was sticky and soft. When the experiment was conducted after 168 h, the rats exhibited mental malaise; they yawned and dozed from time to time; their hair was haggard and erect, showing a light ivory color; the color of the lips, nose, and tail was pale and cyan; grasping resistance was weak; and they had loose stool.

#### 2. Changes in mechanical sensitivity, weight, heart rate and blood pressure

Mechanical allodynia was measured as a reduction compared with the control group in the response threshold to mechanical stimuli applied by von Frey filaments. The neck-skin response thresholds of the CS group and the YCS group were significantly decreased on day 30 after surgery (*P* < 0.01). During sleep deprivation, the response threshold of the rats to mechanical stimuli decreased insignificantly in the control group and CS group but continuously significantly decreased in the YCS group at 72 h (*P* < 0.05), 120 h (*P* < 0.01), and 168 h (*P* < 0.01) (Figure 3a).

Before sleep deprivation, there was no significant difference in body weight among the three groups (p > 0.05). During sleep deprivation intervention, the body weights of the animals increased significantly in the control group and the CS group. By contrast, the body weights of the rats in the YCS group only slightly increased at 72 to 168 h of induction, which might be related to the induction treatments. At the end of the induction period, an increase in weight was obvious in the control group and CS group and was less obvious in the YCS group (*P* < 0.01) (Figure 3b).

Compared with the rats in the control group at the beginning of sleep deprivation, the heart rate and mean blood pressure levels changed insignificantly in each model group. At 168 h of sleep deprivation, the heart rate and mean blood pressure blood pressure of the animals decreased significantly in the YCS group (*P* < 0.01) and changed insignificantly in the control group and CS group. Notably, the heart rates of the rat in the YCS group significantly increased at 0-72 h of sleep deprivation (*P* < 0.01) (Figure 3c). During the entire sleep deprivation process, the mean blood pressure blood pressure of the rats decreased continuously. However, the rats’ heart rate increased from 0 to 72 h and then decreased continuously from 72 to 168 h (Figure 3d).

#### 3. Changes in cAMP, cGMP, and cAMP/cGMP Levels

Before sleep deprivation, there was no significant difference in cAMP, cGMP, or cAMP/cGMP levels among the three groups (*P* > 0.05). After 72 h of sleep deprivation, compared with the control group and the CS group, the serum cAMP of the YCS group was significantly higher (*P* < 0.01), the cGMP content in the serum of the YCS group was significantly lower (*P* < 0.01), and the cAMP/cGMP in the serum of the model group was significantly higher (*P* < 0.01). After 120 h of sleep deprivation, compared with the control group and the CS group, the serum cAMP of the YCS group was significantly higher (*P* < 0.01), and the cAMP/cGMP in the serum of the model group was significantly higher (*P* < 0.05). After 168 h of sleep deprivation, compared with the control group and the CS group, the serum cGMP of the YCS group was significantly higher (*P* < 0.01), and the cAMP/cGMP in the serum of the model group was significantly lower (*P* < 0.01) (Figure 4).

**Figure 4.**
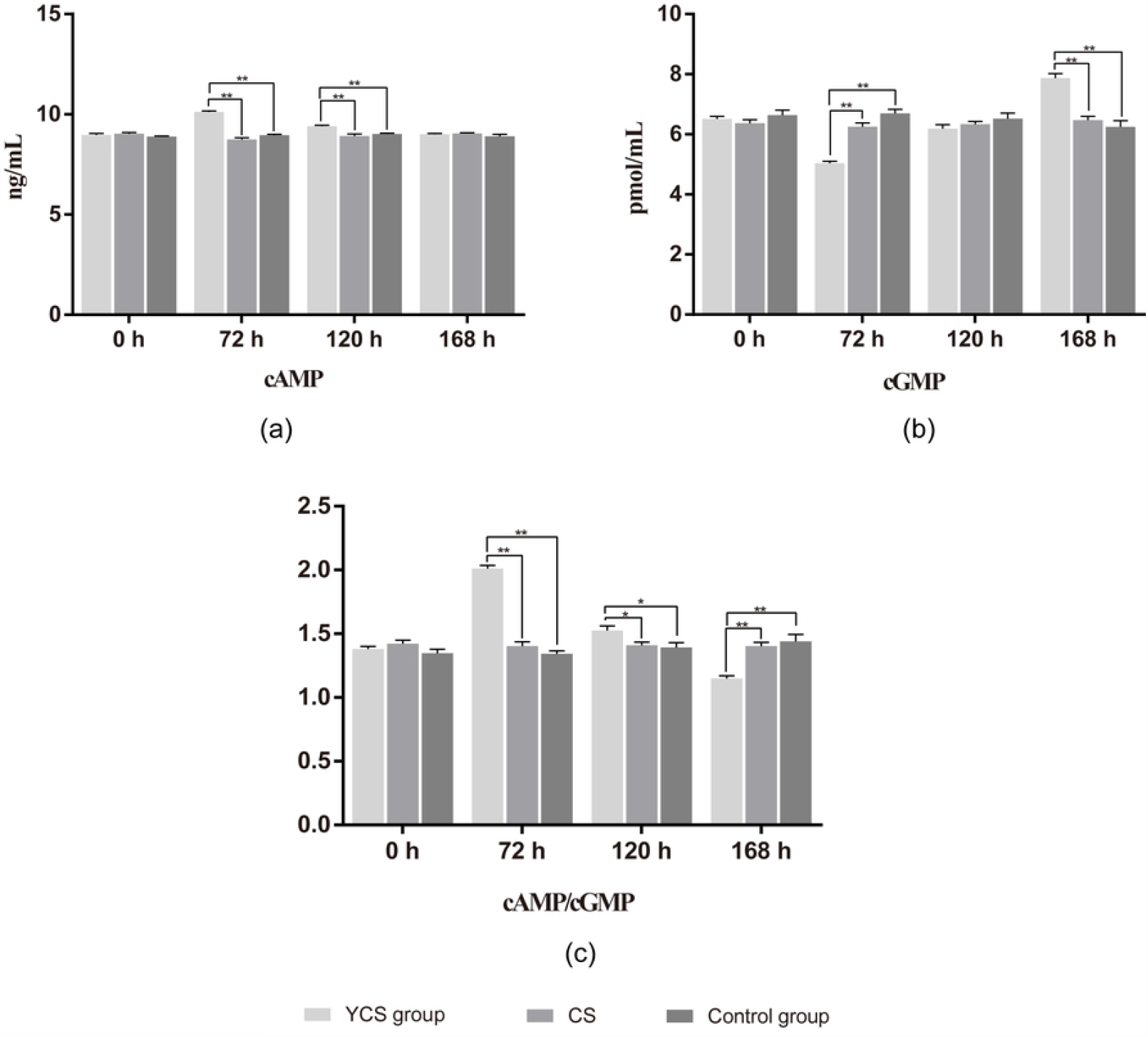
The levels of cAMP, cGMP, and cAMP/cGMP in serum. (a) Changes in cAMP levels of each group measured for 168 hours of sleep deprivation. (b) Changes in cGMP levels of each group measured for 168 hours of sleep deprivation. (c) Changes in cAMP/cGMP levels of each group measured for 168 hours of sleep deprivation. Data are represented as mean ± SEM (n = 9), \**P* < 0.05, \*\**P* < 0.01.

### Indexes of intervertebral disc degeneration in animal models

#### 1. Microstructural changes of the intertebral disk under a light microscope

The collected rat cervical intervertebral discs were stained with H&E and observed at 100 × magnification (Figure 1). As shown in Figure 1c, the intervertebral disc of the control group featured a regular arrangement of the annulus fibrosus and nucleus pulposus and clear cartilage endplate. The intervertebral discs of the CS and YCS groups changed significantly, with the derangement of annulus fibrosus and obvious shrunken nucleus pulposus, and the cartilage endplate exhibited a rough border.

We compared the Miyamoto grading value between the groups by the degree of cervical intervertebral degeneration. As shown in Figure 1d, the value of the control group was significantly lower than that of the other groups (*P* < 0.01). However, there was no significant difference between the CS group and YCS group (after 168 h of sleep deprivation).

#### 2. Ultrastructure of the nucleus pulposus under a transmission electron microscope

As indicated in Figure 2, we observed the ultrastructure of the nucleus pulposus in the different groups. The cells in the control group were round and oval. The cytomembrane was visualized, and the nucleus was in the center with a smooth and integral nuclear membrane. In nucleus was evenly distributed, and heterochromatin was not obvious. Moreover, abundant cellular organs, including rough endoplasmic reticulum, mitochondria, matrix vesicles, and halo, were visible in the cytoplasm (Figure 2c). The cells in the CS group were irregular in shape, and the nuclear membrane structure was uneven. We observed increased heterochromatin, shrinkage of the cytoplasm, decreased organelles, rough endoplasmic reticulum, swollen mitochondria, and thick periplasm (Figure 2b). In the YCS group (after 168 h of sleep deprivation), the cells were similar to those of the CS group (Figure 2a).

#### 3. Protein levels of type II collagen by western blotting

As shown in Figure 5, the level of type II collagen, which is negatively correlated with intervertebral disc degeneration, was higher in the control group than in the other groups (*P* < 0.01). However, there was no significant difference in type II collagen level between the CS group and the YCS group (*P* > 0.05) (Figure 5b).

**Figure 5.**
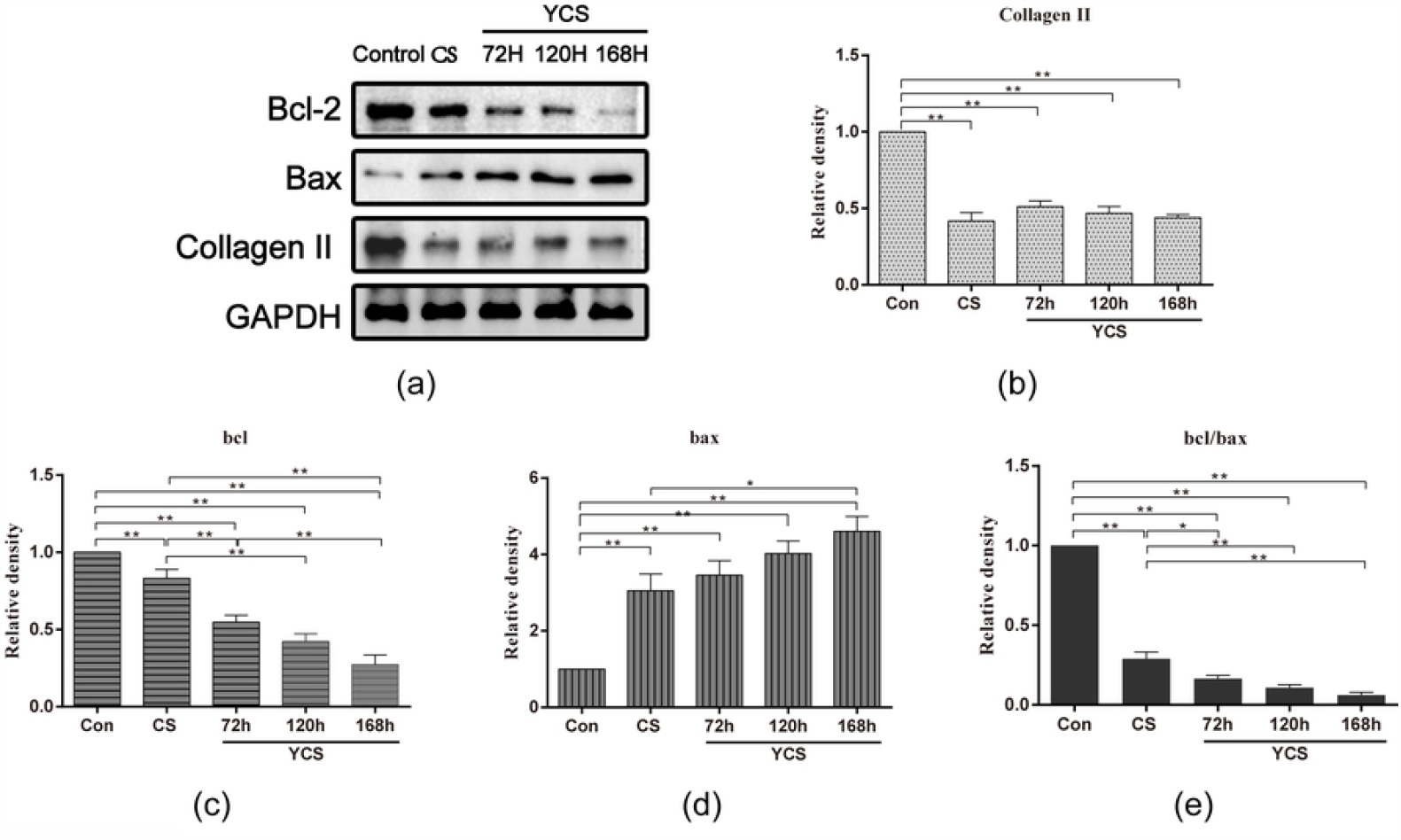
The related protein expression levels of intervertebral disc in animal models. (a) The representative western blotting bands of type II collagen, Bcl-2 Bax and Bcl-2/Bax protein. (b) The protein expression levels of type II collagen. (c) The protein expression levels of Bcl-2. (d) The protein expression levels of Bax. (e) The expression levels of Bcl-2/Bax. Data are represented as mean ± SEM (n = 4), \**P* <0.05, \*\**P* < 0.01.

#### 4. Protein levels of Bcl-2, Bax and Bcl-2/Bax by western blotting

As shown in Figure 5, the level of Bcl-2, which is negatively correlated with apoptosis, was higher in the control group than in the CS groups and the YCS groups (*P* < 0.01). Its level in the YCS group at 72, 120, and 168 h decreased continuously and was lower than that in the CS group (*P* < 0.01) (Figure 5c).

By contrast, the level of Bax, which is positively correlated with apoptosis, was lower in the control group than in the CS groups and the YCS groups (*P* < 0.01). Notably, its level in the YCS group at 72, 120, and 168 h increased continuously and was higher than that in the CS group (Figure 5d).

The level of Bcl-2/Bax was higher in the control group than in the CS groups and the YCS groups (*P* < 0.01). Its level in the YCS group at 72, 120, and 168 h decreased continuously and was lower than that in the CS group (*P* < 0.05) (Figure 5e).

## Discussion

The idea of combining diseases and syndromes of TCM has been used to study CS animal model induction [11]. CS is a disease based on degenerative pathological changes of the intervertebral disc [12]. According to the pathogenesis of a dynamic and static imbalance of CS, the rat CS model is induced by cutting and separating the posterior cervical muscles and ligaments [13]. The induction method is relatively mature and highly reproducible and is widely used in CS research. Our research group also verified the accuracy and feasibility of this model in previous studies [7].

Compared with the control group, the Miyamoto score of the CS group decreased significantly, apoptosis increased, and the mechanical pain threshold decreased significantly. However, the induction method of CS combined with a specific TCM syndrome type has been rarely studied.

Yin deficiency is a specific term in TCM. TCM puts forth the concept that insufficient rest, overwork, and long-term illness will lead to symptoms of Yin deficiency. Modern clinical research has demonstrated that neck pain in patients with cervical spondylopathy with Yin deficiency syndrome is often more severe than other types of CS [14]. Yin deficiency syndrome causes the conditions of CS to become complex, entangling, and difficult to cure. However, the clinical effect of nourishing Yin and tranquilization with TCM is remarkable [15], according to the pathogenesis of Yin deficiency syndrome. At present, there are several published Chinese studies on the induction of Yin deficiency, both of which used the sleep deprivation method to simulate Yin deficiency syndrome. However, this was the first attempt to explore CS complicated with Yin deficiency syndrome [16-20].

The evaluation criteria of the Yin deficiency syndrome model are multidimensional, including behavioral observation, relative biomarkers and conjecture, according to the creation factors that have been proposed in China [12]. Our results showed that compared with the control group and the CS group, the YCS group’s weight decreased significantly at 72, 120, and 168 h of sleep deprivation. Before sleep deprivation intervention, behavioral characteristics of rats in the YCS group were normal. After 72 h of sleep deprivation, the YCS group showed certain fidgety, fluid consumption and internal heat. At this time, the behavior of the YCS group was consistent with the clinical symptoms of Yin deficiency. Moreover, with the extension of sleep deprivation time, rats gradually changed from simple Yin deficiency symptoms to Yin–Yang deficiency symptoms.

Pulse condition refers to the speed, strength, and depth of the pulse. It is an important basis of TCM syndrome differentiation [21]. Yin deficiency syndrome often manifests in a thready and rapid pulse [22]. Because the rat model cannot be accurately measured, our research group plans to use the measured values of rat heart rate and blood pressure to speculate on the changes of rat pulse. An increase in heart rate in the rats indicates a rapid pulse, whereas a decrease in blood pressure means that the pulse is thready and weak. The data measured in our experiment are consistent with the pulse manifestations of Yin deficiency symptoms in rats. At 72 h of sleep deprivation, the heart rate increased and blood pressure decreased, and the heart rate and blood pressure continued to decrease thereafter.

cAMP and cGMP are important second messengers in organisms, and they are antagonistic and participate in the regulation of cell function. This bidirectional regulation is similar to the waxing and waning of Yin–Yang in the theory of Yin and Yang in TCM [23]. Many studies have also confirmed this point. In 1973, the American biologist Goldberg put forward the Yin–Yang hypothesis of biological regulation based on cAMP and cGMP [24]. TCM research found that the cAMP of patients with Yin deficiency was higher and the cAMP/cGMP ratio increased significantly, whereas the cAMP/cGMP ratio decreased significantly in patients with Yang deficiency [25-26]. The results of the ELISA showed that after 72 h of sleep deprivation, the sera cAMP and cAMP/cGMP ratio of the YCS group increased significantly. After 168 h of sleep deprivation, the sera cAMP and cAMP/cGMP ratio of the YCS group decreased. This shows that when sleep deprivation reaches 72 h, the rats exhibit Yin deficiency symptoms, and when it reaches 168 h, there are not only Yin deficiency symptoms but also Yang deficiency symptoms, which is consistent with the symptoms and signs exhibited.

Neck pain is one of the most common symptoms of CS, and it is an important indication for judging the severity of CS. It has been shown that Yin deficiency syndrome exacerbated hyperalgesia in rats with CS. At the same time, hyperalgesia in the YCS group exacerbated gradually with the time of sleep deprivation, which was consistent with the intense pain of patients with CS with Yin deficiency syndrome at night. Furthermore, the apoptosis of intervertebral disc cells in the YCS group is more serious than that in the CS group. However, the change in type II collagen in the intervertebral disc was not obvious in the two groups, indicating that Yin deficiency syndrome exacerbated the apoptosis of intervertebral disc cells.

In this study, we compared models of cervical spondylopathy with Yin deficiency syndrome and cervical spondylopathy in terms of behavior, heart rate and blood pressure, mechanical pain threshold, apoptosis, and disc degeneration. As expected, our results were close to those observed in clinical practice and in human diseases. Thus, our models could be used to explore the pathogenesis of cervical spondylopathy and could serve as an experimental basis of pharmacodynamics and therapeutic research.

## Conclusions

We established rat models of CS with Yin deficiency syndrome, which presented the clinical characteristics specified in TCM. Thus, our models could be used in studies investigating CS with Yin deficiency syndrome.

## Supporting information

**S1 Dataset. All data obtained during this study**.

(XLSX)

**S1 Raw images**.

(DOCX)

## Acknowledgments

We extend our gratitude to the animal care staff of the Animal Experiment Center of First Affiliated Hospital of Shandong First Medical University & Shandong Provincial Qianfoshan Hospital for their support and assistance in the maintenance and feeding of rats. We also thank all the participants of this study.

## Authors’ Contributions

**Conceptualization:** Wenlong Yang, Fangming Liu

**Data curation:** Wenlong Yang, Muqing Liu.

**Formal analysis:** Wenlong Yang, Qinran Sun, Wenqing Wu.

**Funding acquisition:** Wenlong Yang, Fangming Liu.

**Investigation:** Peng Gao, Zhizhen Liu.

**Methodology:** Muqing Liu, Peng Gao

**Project** administration: Lei Liu, Fangming Liu.

**Resources:** Muqing Liu, Zhizhen Liu. **Software:** Wenlong Yang, Zhizhen Liu. **Visualization:** Qinran Sun, Wenqing Wu.

**Writing - original draft:** Wenlong Yang, Zhizhen Liu.

**Writing - review & editing:** Lei Liu, Fangming Liu.

